# Screening of Oyster Peptides for Anti-Muscle Atrophy Based on Machine Learning and Computer Simulation: Guided by Antioxidant Pathways

**DOI:** 10.1101/2025.10.28.685008

**Authors:** Jiyuan Zhang, Lunzhou Lin, Jiawei Yang, Hailin Zheng, Ting Fang, Wenjuan Li

## Abstract

Muscle atrophy poses a serious threat to human health, with its primary pathogenic mechanisms closely linked to oxidative stress. This study focuses on the potential of oyster peptides in alleviating dexamethasone (DEX)-induced skeletal muscle atrophy and their underlying antioxidant mechanisms. Utilizing efficient integrated machine learning and computer simulation methods, a systematic screening of active peptides and mechanistic research was conducted. The results revealed that oyster peptides at concentrations of 25–50 μg/mL significantly improved the decline in cell viability and myotube atrophy induced by DEX, while downregulating the expression of muscle atrophy markers *Atrogin-1* and *MuRF1*. Through LC-MS/MS, 220 high-activity peptide sequences were identified. Following the replication and extension of the iAnOxPep integrated learning model, 15 potential antioxidant peptides were selected. Among them, the short peptide AWPGPQ demonstrated the strongest binding affinity with Keap1 and PPARγ targets. Molecular dynamics simulations confirmed its stability, suggesting that AWPGPQ may exert its dual effects—antioxidant and anti-muscle atrophy—by modulating the *Keap1-Nrf2* and *PPARγ* signaling pathways. This study established a systematic and efficient strategy for screening natural active peptides, providing theoretical support and technical pathways for the discovery of multifunctional short peptide candidates, with significant theoretical value and application prospects.

## 1. Introduction

Muscle atrophy is a pathological process characterized by a reduction in the cross-sectional area of muscle fibers and a decrease in the number of muscle cells [1], leading to the progressive loss of muscle mass and strength [2]. In the complex mechanisms underlying muscle atrophy, oxidative stress has been confirmed by several studies to play a key role [3]. Research by Mukai and Terao [4] demonstrated that oxidative stress promotes muscle protein degradation by activating ubiquitin ligases such as *atrogin-1* and *MuRF-1* [5], thereby accelerating the atrophy process. The research team of Marco Segatto [6] targeted and regulated ROS metabolic regulators, closely linked to oxidative stress, such as BET proteins, and significantly improved the muscle atrophy phenotype in a Duchenne muscular dystrophy (DMD) mouse model, further confirming the critical role of oxidative stress pathways in the occurrence and development of muscle atrophy.

Antioxidant interventions have been regarded as a potential strategy for alleviating muscle atrophy. However, traditional synthetic antioxidants (such as phenolic compounds) suffer from shortcomings like poor thermal stability and the potential to generate toxic quinone by-products [7], which limits their clinical application. This has driven researchers to focus on the development of efficient, low-toxicity natural antioxidants, with food-derived bioactive peptides gaining attention due to their small molecular weight, strong targeting ability, and high safety [8]. Hur *et al*. [9] confirmed that beef peptides can alleviate DEX-induced myotube atrophy by upregulating *SOD* expression. Oyster peptides have also been shown by Zhu and Xiang’s research team to exhibit significant antioxidant activity and immune-modulatory functions [10,11], indicating their promising potential for combating muscle atrophy. Notably, the Böttcher team [12] confirmed that the activation of the *Keap1-Nrf2* pathway effectively alleviates oxidative stress-induced muscle degeneration, while the Neha Rani team [13] demonstrated that the activation of the *PPARγ* pathway can significantly inhibit oxidative stress, thereby improving muscle damage. Research by Dovinova *et al*. [14] also revealed the synergistic antioxidant effects of the *Keap1-Nrf2* and *PPARγ* pathways, yet studies on how natural active peptides specifically regulate this synergistic network remain insufficient.

Therefore, current research still faces three major limitations: (1) unclear active components, often remaining at the crude extract level, lacking precise identification of core active peptide sequences [15,16]; (2) insufficient mechanistic explanations, especially regarding how oyster peptides regulate multiple pathways, such as *Keap1-Nrf2* and *PPARγ*, remains unexplored [17]; (3) low screening efficiency, relying on low-throughput empirical screening [18]. To overcome these bottlenecks, there is a pressing need to integrate advanced computer technologies such as machine learning to establish an efficient and precise active peptide screening and mechanism research system.

Based on this, the present study aims to establish a system that integrates computational and experimental biology for systematic screening, to efficiently identify oyster peptides with significant dual functions in antioxidant and anti-muscle atrophy activities, and to explore their potential biological mechanisms. Methodologically, we will make full use of machine learning and computer simulation technologies [19] and have developed a systematic screening and mechanistic research strategy that integrates LC-MS/MS peptide identification, integrated machine learning model prediction [20,21], multi-target molecular docking [22,23], molecular dynamics simulation validation [24], and ADMET drugability evaluation. Through this innovative screening strategy, we aim not only to identify effective oyster peptides with dual functions but also to delve deeper into their molecular mechanisms of resisting muscle atrophy through multi-pathway synergistic actions. Furthermore, the research outcomes will provide new natural-source intervention strategies for the prevention and treatment of muscle atrophy [25], and will significantly promote the deep development and application of oyster peptides in the fields of biomedicine and functional foods.

## 2. Results

### 2.1. Molecular Weight Distribution and Basic Nutritional Composition of Oyster Peptides

Oyster peptides were prepared by alkaline protease hydrolysis, and their physicochemical properties were characterized through molecular weight distribution and basic composition analysis. The molecular weight (MW) distribution is an important indicator for assessing the degree of protein hydrolysis and guiding subsequent separation and purification. The results indicated that the enzymatic hydrolysis products were predominantly low molecular weight peptides, with components below 3 kDa accounting for 71.94%. Specifically, the proportions of peptides in the 180-500 Da, 500-1000 Da, 1000-2000 Da, and 2000-3000 Da ranges were 6.93%, 15.31%, 11.87%, and 36.91%, respectively. Basic composition analysis (Table 1) showed that oyster peptides are characterized by high protein content, low sugar, and low moisture. These compositional features not only indicate a high concentration of bioactive peptides but also contribute to the product’s excellent storage stability, confirming oysters as a valuable marine protein resource with great development potential.

**Table 1.** Basic composition of OP.

| Component | Dry basis percentage |
| --- | --- |
| Protein | 81.3 ± 0.12% |
| Total Sugar | 2.78 ± 0.18% |
| Moisture | 10.22 ± 0.07% |

### 2.2. The Effect of Oyster Peptides on C2C12 Cell Viability and DEX-Induced Damage

As shown in Figure 1a, 24-hour treatment with 5 μM DEX did not significantly affect C2C12 cell viability, while 10, 20, and 40 μM DEX treatments significantly reduced cell viability. Specifically, 10 μM DEX caused a 27.98% decrease in cell viability compared to the control group. Based on this, 10 μM was selected as the concentration for inducing muscle atrophy in subsequent models.

**Figure 1.**
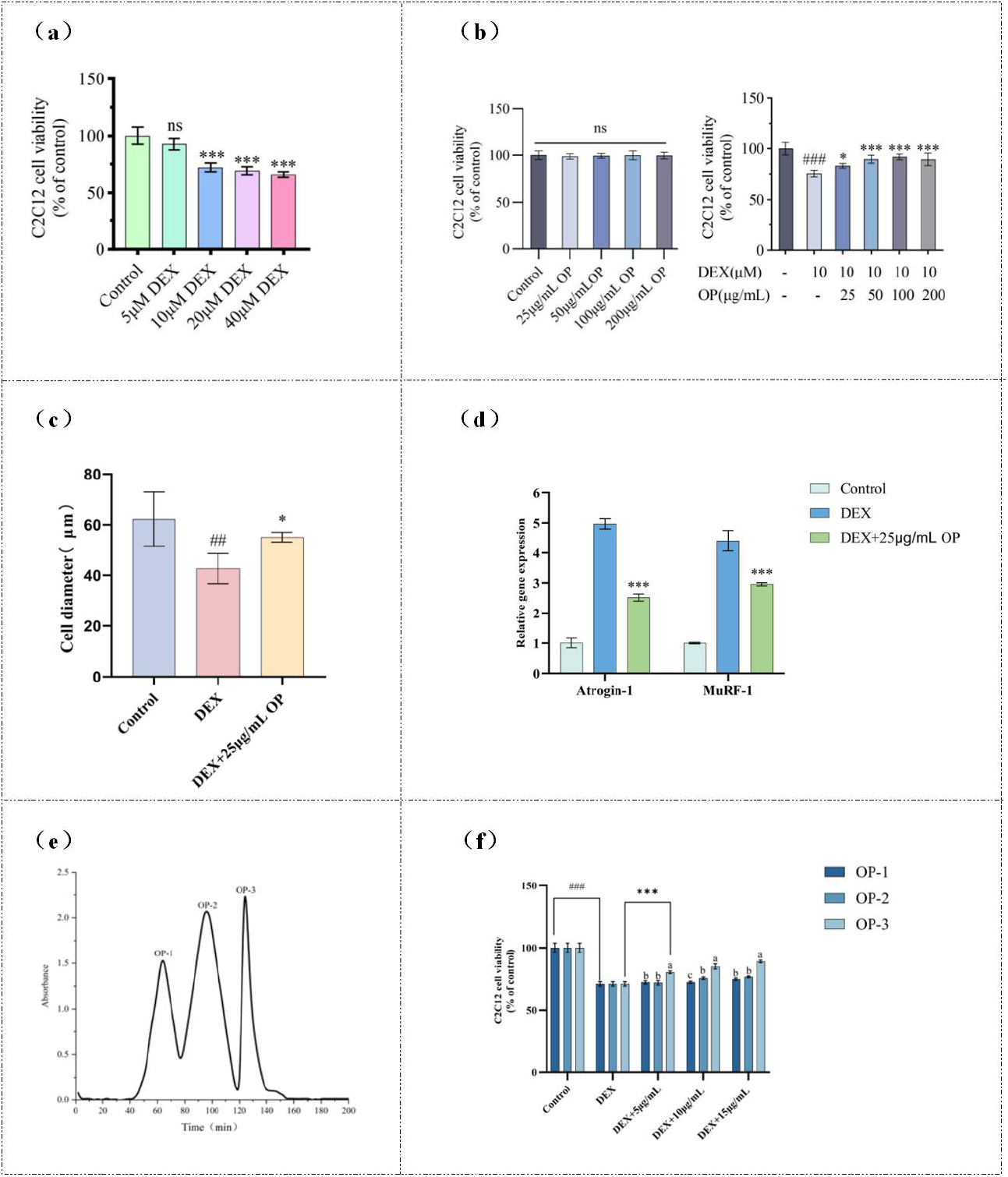
Effects of Oyster Peptides (OP) on DEX-Induced C2C12 Muscle Atrophy Model and Bioactive Component Screening:(a) Effects of different concentrations of DEX on C2C12 cell viability;(b) Cytotoxicity assessment of OP on C2C12 cells and its protective effect against DEX-induced cell viability reduction;(c) Improvement of myotube diameter by OP treatment;(d) Regulation of muscle atrophy-related genes *Atrogin-1* and *MuRF1* by OP;(e) Chromatographic profile of OP fractions separated by G-25 gel filtration;(f) Effects of OP fractions (OP-1, OP-2, OP-3) on C2C12 cell viability.

Treatment with OP (25, 50, 100, 200 μg/mL) alone showed no significant toxicity to C2C12 cells (Figure 1b), with cell viability remaining similar to that of the control group. Further intervention with varying concentrations of OP in the 10 μM DEX-induced model significantly improved the reduction in cell viability induced by DEX. The 25 μg/mL OP group showed a 7.43% increase in cell viability compared to the model group. As the concentration increased, cell viability also improved, although it plateaued at 50 μg/mL and did not increase further with higher concentrations.

### 2.3. The Effect of Oyster Peptides on Myotube Diameter

To visually observe the improvement in myotube atrophy by OP, immunofluorescence experiments were conducted to detect MyHC protein levels, and myotube diameters were quantitatively analyzed. In the DEX group, MyHC protein fluorescence expression was significantly lower than in the control group, indicating muscle atrophy. However, in the DEX + 25 μg/mL OP group, MyHC expression was significantly higher than in the DEX group, indicating that atrophy was alleviated (Figure 2). The myotube diameter in the DEX group was significantly reduced, while OP treatment significantly restored the myotube diameter (Figure 1c). In summary, OP effectively improved DEX-induced myotube atrophy in C2C12 cells.

**Figure 2.**
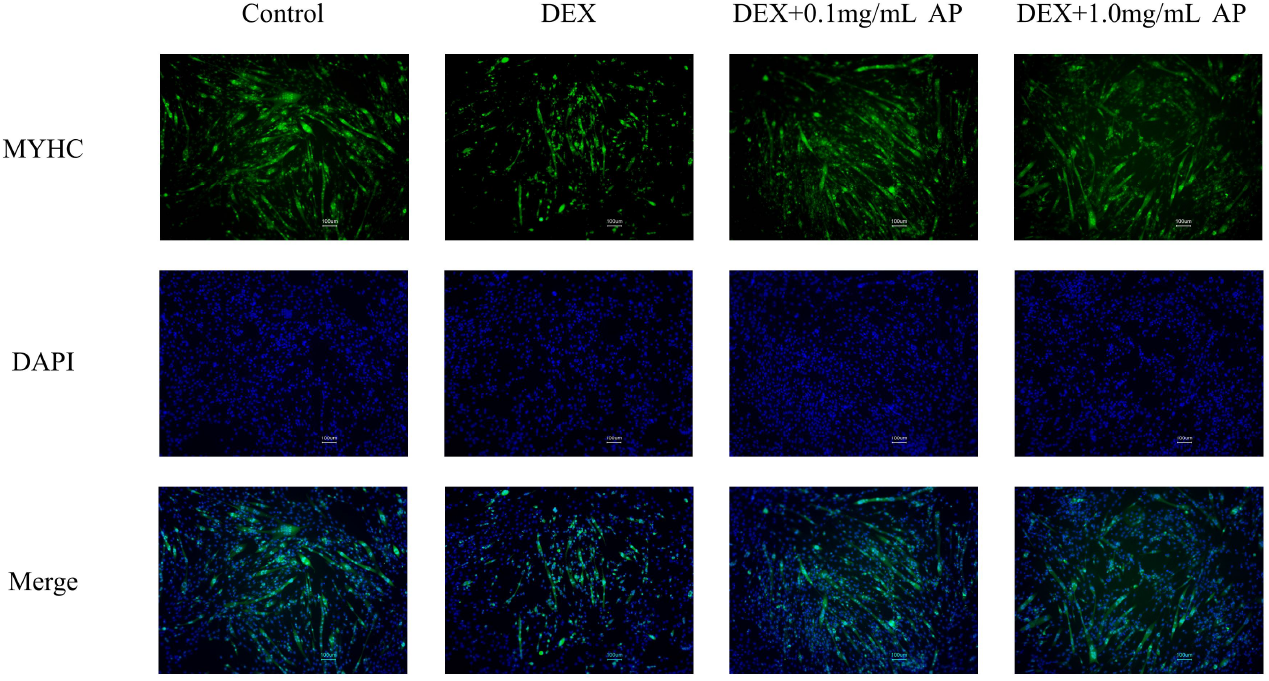
Immunofluorescence staining of C2C12 myotubes in the DEX group and the OP treatment group

### 2.4. Regulation of Muscle Atrophy-Related Gene Expression by Oyster Peptides

The UPS system plays a key role in regulating cellular protein degradation, with Atrogin-1 and MuRF1 being core ubiquitin ligases upregulated by oxidative stress. Their expression levels are commonly used as important markers of muscle atrophy [26]. Chen et al. [27] pointed out that oxidative stress can activate the FoxO transcription factor pathway, inducing the expression of Atrogin-1 and MuRF1, thereby promoting muscle protein degradation and muscle atrophy. Compared to the control group (Figure 1d), the expression of Atrogin-1 and MuRF1 genes was significantly upregulated in the DEX model group (P<0.001). After OP intervention, the expression levels of Atrogin-1 (P<0.001) and MuRF1 (P<0.001) were reversed, further indicating that OP can effectively improve muscle atrophy caused by oxidative stress induced by DEX.

### 2.5. LC-MS/MS Identification of High-Activity Peptide Sequences

To identify the core bioactive peptide sequences in oyster peptides, three components (OP-1, OP-2, OP-3) obtained from initial purification using Sephadex G-25 gel chromatography were compared. In the cell viability assay, the OP-3 fraction exhibited significant anti-muscle atrophy activity, thus OP-3 was selected for further in-depth analysis (Figure 1e, Figure 1f).

After reduction and alkylation treatment (DTT reduction and IAA alkylation), the OP-3 fraction was subjected to LC-MS/MS analysis using the Easy-nLC 1200 high-performance liquid chromatography system coupled to the Orbitrap Fusion Lumos high-resolution mass spectrometry platform. A total of 9,630 scan records were obtained, and 44,310 raw peptide sequences were identified.

To further improve the reliability of peptide identification, PEAKS software was used for rigorous filtering of the mass spectrometry data. Based on a false discovery rate (FDR) control of 1.0% and combining a De novo scoring mechanism, only peptides with a De novo score greater than 90% were retained. A total of 603 high-quality, reliable peptide sequences were obtained. These 603 peptides were further evaluated for their potential biological activity using the PeptideRanker online prediction platform, which assigned a biological activity prediction score ranging from 0 to 1 to each peptide. According to the prediction results, 220 peptides scored ≥ 0.7, indicating a strong potential for biological activity and warranting further investigation for their potential as anti-muscle atrophy functional peptides.

### 2.6. Integrated Machine Learning Prediction of Potential Antioxidant Peptides

To ensure the reliability of the prediction results, this study employed five evaluation metrics—Accuracy, AUC, MCC, Specificity, and Sensitivity—to comprehensively assess a total of 98 machine learning base models using the bootstrap method. As shown in Figure 3d, the newly introduced support vector machine (SVM) and multilayer perceptron (MLP) models performed excellently across all metrics, outperforming most of the classifiers in the original model. In particular, the SVM classifier showed outstanding performance in Specificity, with scores exceeding 0.85 for features such as AAC, PC-PAAC, and SC-PAAC. The LGBM classifier maintained stable performance, with an accuracy and sensitivity exceeding 0.80, and an AUC value close to 0.85, indicating good classification capability. Although the ETC and RF models showed slightly weaker performance with KSDC and DPC features, they still exhibited strong predictive ability with PDT and SC-PAAC.

**Figure 3.**
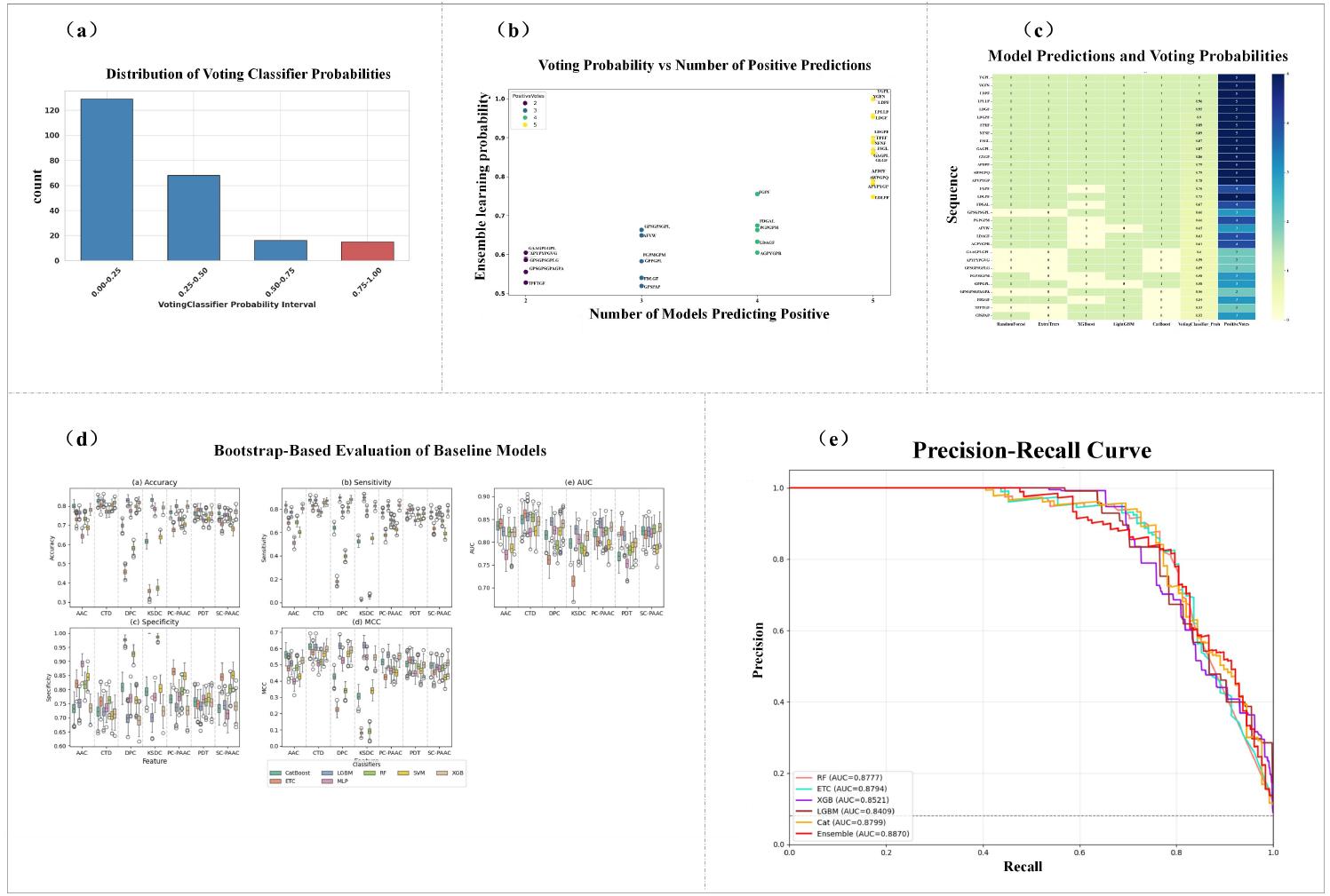
Voting Results and Performance Evaluation of the Ensemble Model: (a) Probability interval distribution of ensemble voting classifier outputs, showing the number of samples within each prediction interval; (b) Scatter plot of predicted probability versus number of base models predicting a positive label, revealing the degree of consensus among classifiers; (c) Heatmap of prediction results from each base model and the ensemble voting classifier, illustrating consistency and variability across models; (d) Distribution of multiple performance metrics—Accuracy (ACC), Area Under the Curve (AUC), Matthews Correlation Coefficient (MCC), Sensitivity, and Specificity—across 100 bootstrap resampling iterations for each classifier; (e) Precision-Recall (PR) curves of multiple models, comparing classification performance and AUC values of individual base learners and the ensemble model.

The integrated learning method effectively enhanced the model’s generalization ability by combining the strengths of multiple base learners. The evaluation results demonstrated that the integrated model built using 98 feature vectors outperformed any individual model in terms of accuracy, AUC, and other core metrics, and performed stably in the Precision-Recall curve, particularly in the upper-right region, demonstrating stronger overall performance (Figure 3e). Compared to the iAnOxPep model proposed by Mir Tanveerul Hassan et al., the AUC value of the integrated model in this study improved from 0.8673 to 0.8870, marking a 2.27% performance enhancement. Overall, the integrated model improved overall prediction performance by merging multiple base learners and optimizing strategies such as dataset expansion, feature standardization, and automated hyperparameter search, leading to improved balance between precision and recall over the iAnOxPep model.

The integrated learning model was then used to predict the 220 high-confidence, high-bioactivity peptide sequences, and the predicted probabilities were divided into four intervals with a 0.25 interval. The 0.50–0.75 and 0.75–1.00 intervals contained 16 and 15 peptides, respectively, accounting for a total of 13.6% (Figure 3a). This indicates that the number of peptides with high predicted probabilities is relatively low, showing strong screening specificity. To comprehensively analyze the prediction results from both base learners and the integrated model, we set a threshold of 0.5 for positive predictions and visualized the scatter plot and heatmap of the 31 peptides predicted as positive labels by the integrated learning model. The integrated model’s prediction probability (Ensemble learning probability) increased significantly with the number of models predicting positive labels (Number of Models Predicting Positive), particularly when the prediction probability exceeded 0.75, where nearly all models predicted positive labels consistently, reflecting high consistency between the integrated model and its sub-models (Figure 3b, Figure 3c).

Finally, 15 peptides with a prediction probability greater than 0.75 (including AWPGPQ, NFNF, FGPF, APYPYGP, LDGPF, YGPL, YGFN, APDPF, LDPF, FPEF, FSGL, LDGF, GLGF, GAGPL, LPLLP) were selected as high-potential antioxidant peptides and prioritized for subsequent validation and experimental studies.

### 2.7. Molecular Docking Validation of the Affinity Between Dual-Function Peptides and Their Targets

The antioxidant peptides predicted from machine learning were used as ligands in molecular docking with target receptors to further validate their dual functionality in antioxidant and anti-muscle atrophy activities. Considering the central roles of *Keap1* and *PPARγ* in regulating oxidative stress and muscle atrophy pathways, as well as supporting literature evidence, these two receptors were selected as target proteins for docking.

*Keap1* is a core regulatory factor in the oxidative stress response [28], directly involved in oxidative stress by inhibiting *Nrf2*-mediated antioxidant gene expression [29]. The latest research by the Böttcher team further confirmed that activation of the *Nrf2-Keap1* pathway can effectively improve muscle cell degeneration [12]. The Neha Rani team [13] demonstrated that activation of the *PPARγ* pathway significantly inhibits AGE-RAGE-mediated oxidative stress, thereby improving muscle damage. Research by Dovinova et al. [14] also revealed the synergistic antioxidant effects of the *Keap1-Nrf2* and *PPARγ* pathways. Therefore, Keap1 and PPARγ were chosen as ideal target receptors to validate the dual functionality of oyster peptides in antioxidant and anti-atrophy activities.

The 15 antioxidant peptide sequences selected from the machine learning model, along with the YYR sequence and the control substance rosiglitazone (RSG), were docked with the Keap1 and PPARγ receptors.

The results of molecular docking are shown in Table 2. Among the 15 peptide sequences assessed (excluding the control substances RSG and YYR), 6 peptides (AWPGPQ, NFNF, FGPF, APYPYGP, APDPF, YGFN) had binding energies with the *PPARγ* receptor lower than −8 kcal/mol, indicating strong binding affinity for this target, suggesting that they may exert their activity through the *PPARγ* pathway.

**Table 2.**
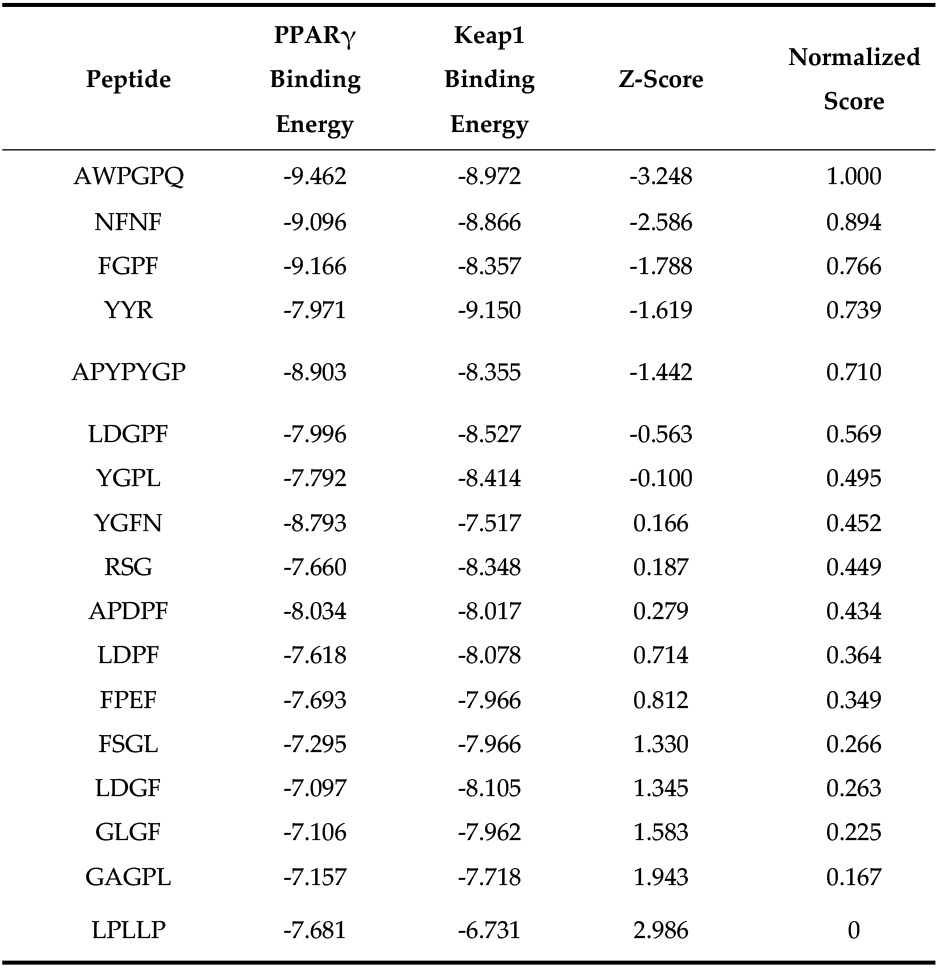
Peptide Binding Energies and Scoring for PPARγ and Keap1.

**Table 3.**
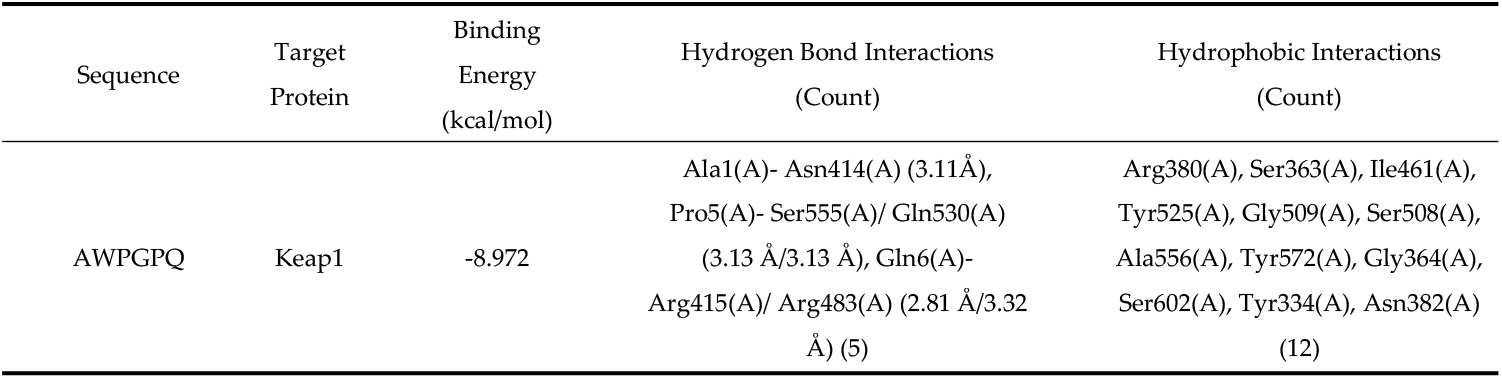
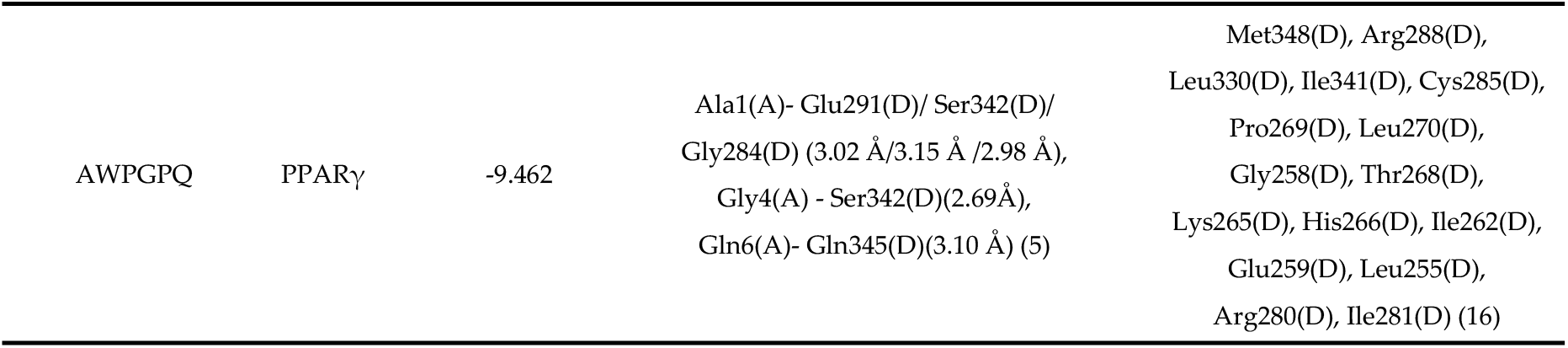
Details of the interactions between AWPGPQ, Keap1 and PPARγ.

Meanwhile, 9 peptides (AWPGPQ, NFNF, FGPF, APYPYGP, LDGPF, YGPL, APDPF, LDPF, LDGF) showed binding energies with Keap1 lower than −8 kcal/mol, implying that they may exert their activity through the *Keap1-Nrf2* pathway. Further analysis of the Z-score revealed that three peptides (AWPGPQ, NFNF, FGPF) had significantly better overall Z-scores compared to the control active substances RSG and YYR, indicating that these peptides may simultaneously and efficiently regulate both the *PPARγ* and *Keap1-Nrf2* pathways, with potential synergistic effects in antioxidant and anti-muscle atrophy activities.

Among the peptides, the sequence AWPGPQ exhibited the best score, with binding energies of −9.462 kcal/mol and −8.972 kcal/mol with the two major receptor proteins, respectively. This peptide was identified as the most promising candidate for further research into its antioxidant and anti-muscle atrophy activity.

### 2.8. Analysis of Hydrogen Bonding and Hydrophobic Interactions Between Peptides and Targets

Molecular visualization of the binding mode between the peptide sequence AWPGPQ and the receptor proteins was performed using Pymol, as shown in Figure 4a. In the binding with Keap1, the peptide forms hydrogen bonds with Ala1(A) and Asn414(A) at a distance of 3.11 Å, Pro5(A) with Ser555(A) and Gln530(A) at distances of 3.13 Å, and Gln6(A) with Arg415(A) and Arg483(A) at distances of 2.81 Å and 3.32 Å, respectively. These hydrogen bonds enhance the affinity and binding strength between the peptide and the target. In the binding with PPARγ, Ala1(A) forms hydrogen bonds with Glu291(D), Ser342(D), and Gly284(D) at distances of 3.02 Å, 3.15 Å, and 2.98 Å, respectively. Gly4(A) forms a hydrogen bond with Ser342(D) at 2.69 Å, and Gln6(A) forms a hydrogen bond with Gln345(D) at 3.10 Å. According to the studies of Wu et al. [30,31], the optimal distance for hydrogen bonds is generally between 2.7 Å and 3.2 Å. Hydrogen bonds within this range significantly increase the specificity and stability of the binding.

**Figure 4.**
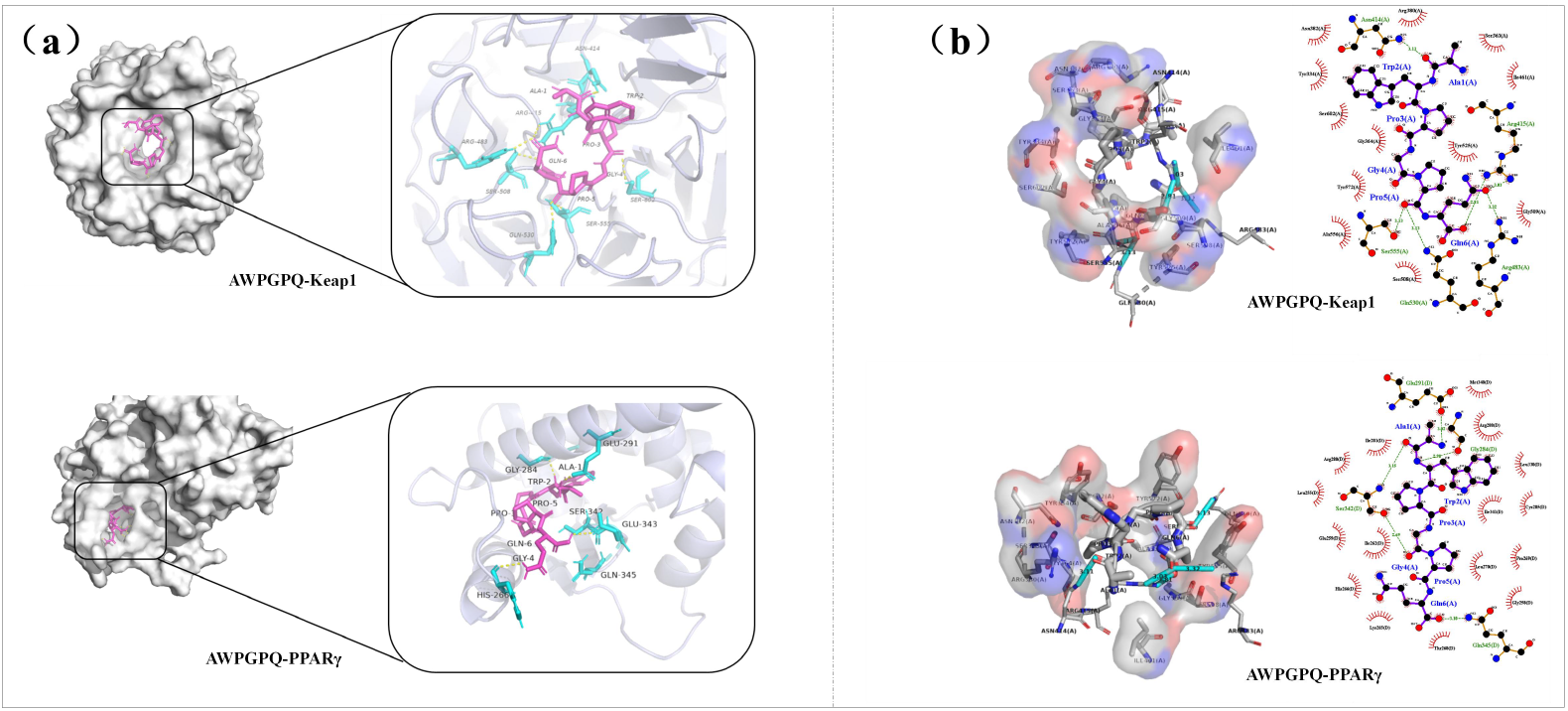
Molecular Docking and Binding Mode Analysis of Peptide AWPGPQ with Keap1 and PPARγ:(a) Binding conformation analysis of AWPGPQ with Keap1 and PPARγ based on PyMOL visualization, showing spatial fitting, hydrogen bonding interactions, and involvement of key residues;(b) Binding mode analysis using LigPlus, illustrating hydrogen bonds and hydrophobic interactions between the peptide and the two receptors, as well as binding characteristics with critical target residues.

Hydrophobic interactions between the target and receptor were further analyzed using Ligplus, as shown in Figure 4b. Hydrophobic regions in the peptide, such as Ala1(A) and Trp2(A), interact with hydrophobic residues in Keap1 such as Arg380(A), Ser363(A), Ile461(A), Tyr525(A), Gly509(A), Ser508(A), Ala556(A), and Tyr572(A), stabilizing the binding through hydrophobic interactions. Similarly, hydrophobic residues in the peptide interact with hydrophobic residues in PPARγ, including Met348(D), Arg288(D), Leu330(D), Ile341(D), and Cys285(D), further reducing water molecule interference and reinforcing the stability of the binding interface.

### 2.9. Molecular Dynamics Simulation to Validate the Stability of Peptide-Target Complexes

Based on the results of molecular docking, the active peptide sequence AWPGPQ exhibits strong binding energy and a tight binding mode with both Keap1 and PPARγ receptor proteins. However, this result alone cannot eliminate potential errors and uncertainties [32], such as the limitations of static docking assumptions and the effects of hydration and solvent interactions on binding [33]. To address this, we performed 80 ns molecular dynamics simulations for each peptide-receptor complex and further assessed the stability of the complexes by analyzing multiple stability indicators and interaction parameters [34].

The complex of AWPGPQ with the Keap1 receptor showed an RMSD (Root Mean Square Deviation) curve that stabilized between 0.1 nm and 0.2 nm during the 80 ns simulation (Figure 5a), indicating that the complex maintained a high structural stability throughout the simulation. The RMSF (Root Mean Square Fluctuation) values showed slight fluctuations at certain residues within the binding pocket (Figure 5c), but remained below 0.3 nm, reflecting minimal fluctuation in this region, which stayed within a stable range. By calculating the probability density of the conformational distribution using RMSD and Rg (radius of gyration), we generated a free energy landscape. This revealed that the binding of Keap1 to AWPGPQ exhibits strong affinity (Figure 5e,f), and the complex remained in a low free energy region (blue zone) during the simulation, indicating that this binding mode was highly conserved throughout the simulation, remaining close to the initial conformation.

**Figure 5.**
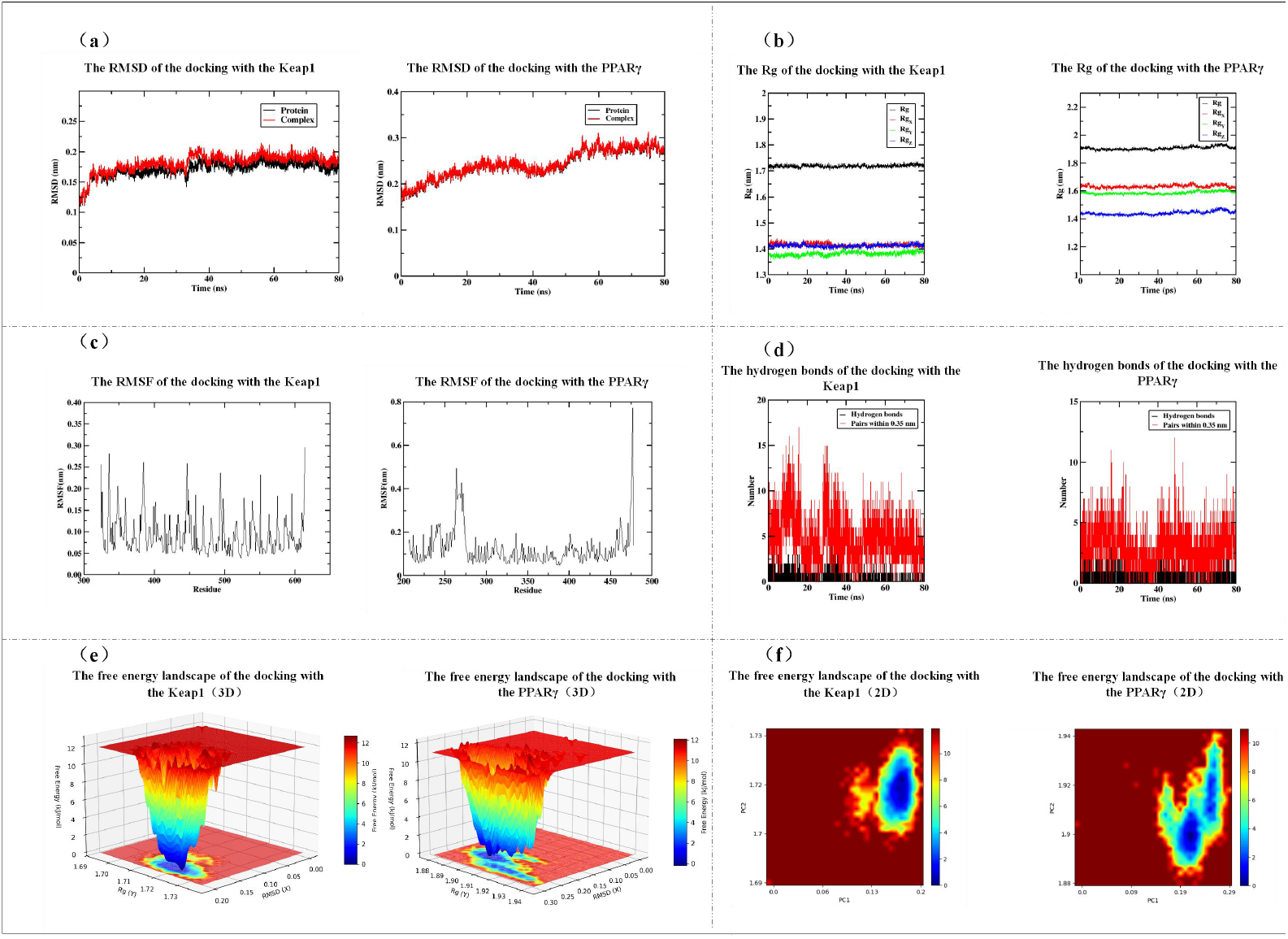
Molecular Dynamics Simulation Analysis of AWPGPQ Complexes with Keap1 and PPARγ Receptors: (a) RMSD (root-mean-square deviation) profiles of Keap1 and PPARγ complexes over the 80 ns simulation period; (b) Evolution of radius of gyration (Rg) for peptide–protein complexes, reflecting compactness of the systems; (c) RMSF (root-mean-square fluctuation) analysis of Keap1 and PPARγ complexes, indicating residue-level flexibility; (d) Variation in the number of hydrogen bonds formed during simulation, demonstrating the interaction stability between the peptide and both receptors; (e) Three-dimensional free energy surface plots of Keap1 and PPARγ complexes; (f) Two-dimensional free energy landscapes derived from principal component analysis (PCA), revealing conformational stability and dominant conformer distributions.

**Figure 6.**
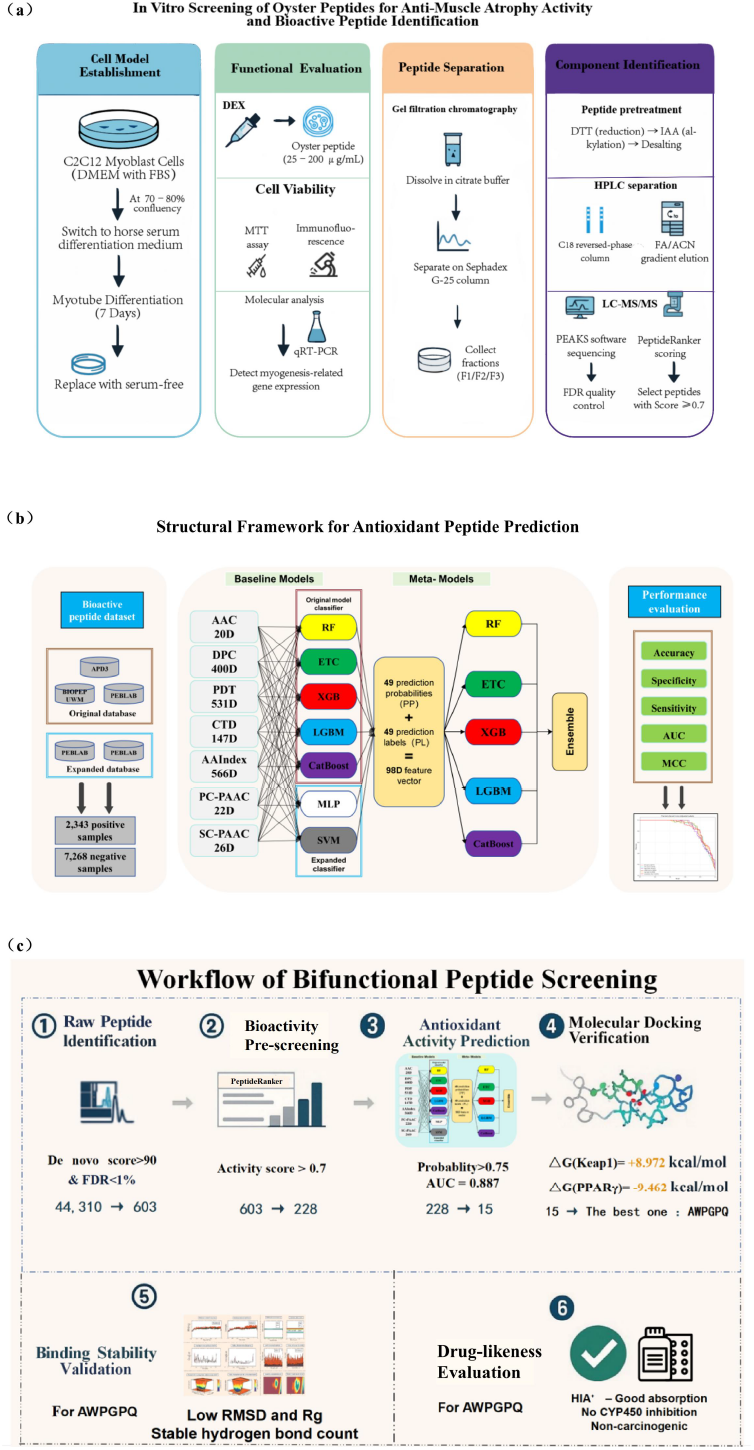
Technical Flow Diagram: (a) In Vitro Screening of Oyster Peptides for Anti-Muscle Atrophy Activity and Bioactive Peptide Identification; (b) Improved iAnOxPep Integrated Model Flowchart: Antioxidant Peptide Prediction System Architecture; (c) Dual-Function Peptide Screening Flowchart.

For the complex of AWPGPQ with the PPARγ receptor, the RMSD value showed a slight increase at the 5 ns mark (Figure 5a), suggesting that the active region of the complex may have undergone local adjustments (Figure 5c, fluctuation of residues 260-275 on the right side). The free energy landscape displayed two closely adjacent blue regions (Figure 5e, 5f), which reflects the transition state process. Despite this, the RMSD remained below 0.3 nm, indicating that the overall fold remained stable, with no significant structural changes. The Rg curve also showed relatively smooth changes (Figure 5b), suggesting that the complex did not undergo significant expansion or deformation during the simulation. The number of hydrogen bonds fluctuated between 1 and 3, indicating a flexible hydrogen bond network in the complex (Figure 5d). This flexible structural network helps maintain the stability of the complex while ensuring a degree of adaptability [35].

These molecular-level evidence collectively suggest that the complex of AWPGPQ with both PPARγ and Keap1 receptors exhibits a tight and stable binding mode, achieving its dual functionality in antioxidant and anti-muscle atrophy through multi-target actions.

### 2.10 ADMET Prediction Analysis

Based on the ADMET prediction results, the active peptide AWPGPQ demonstrated excellent human intestinal absorption (HIA+, probability 0.8825), but low permeability through Caco-2 cells (Caco2-, probability 0.9020), suggesting that its oral bioavailability may need to be enhanced through formulation strategies [36]. The compound shows no carcinogenic risk (probability 0.9212) and no genetic toxicity (Non-AMES toxic, probability 0.9034), confirming its safety advantages. Notably, it exhibits weak inhibition of the hERG potassium channel (Weak inhibitor, probability 0.9525), highlighting the need to monitor cardiac safety in future optimizations. The acute oral toxicity classification is Type III, and the toxicity risk can be further reduced through molecular design. Furthermore, the peptide is unlikely to cross the blood-brain barrier (BBB-, probability 0.9079), which may help avoid central nervous system adverse effects [37]. In terms of metabolism, AWPGPQ does not act as a substrate or inhibitor for any major CYP450 enzymes (including 1A2, 2C9, 2D6, 2C19, and 3A4) (probability >0.72), indicating its excellent metabolic stability and very low risk of drug-drug interactions [38].

## 3. Discussion

This study investigates the potential of oyster peptides in antioxidant and anti-muscle atrophy activities, systematically exploring their biological activity and mechanisms starting from cellular models. The experiments revealed that, within the concentration range of 25 to 50 μg/mL, oyster peptides effectively counteracted DEX-induced C2C12 cell damage, significantly enhancing cell viability and improving myotube morphology. During this process, we observed a notable upregulation of *MyHC* expression, suggesting that oyster peptides may directly participate in the synthesis and maintenance of muscle fiber structural proteins. Further qRT-PCR analysis supported this hypothesis, as it significantly suppressed the expression of muscle atrophy-related *Atrogin-1* and *MuRF1*, indicating that oyster peptides may inhibit protein degradation by intervening in the ubiquitin-proteasome pathway. Compared to existing studies, such as those by Jang *et al*. on fermented whey protein peptides [39] and Lin *et al*. on tilapia peptides [40] in muscle atrophy models, we systematically evaluated the cytoprotective effects of oyster peptides and observed more significant improvements in cell viability and gene expression regulation at similar concentration levels, suggesting that oyster peptides possess a unique composition of biological activity and regulatory mechanisms.

These results not only confirm the intervention potential of oyster peptides in muscle atrophy but also provide preliminary insights into their possible biological effects through specific molecular mechanisms. Building upon these findings, we established a natural peptide screening process integrating mass spectrometry analysis, machine learning prediction, molecular docking, molecular dynamics simulation, and ADMET evaluation. This “sequence-to-activity-to-mechanism-to-safety” closed-loop system not only improves screening efficiency but also lays a technological foundation for future natural functional peptide research. Unlike traditional functional peptide research, such as that by Hao, Wang, and others, which relies on empirical screening and low-throughput validation [16,18], our study centers on a data-driven predictive model, significantly enhancing screening efficiency and targeting mechanism accuracy, particularly suited for high-throughput development of marine peptide resources. Especially in the context of increasing emphasis on targeted mechanisms and drug development potential in natural product research [41], this methodology has broad applicability and significant potential for promotion.

Looking forward, research will continue to focus on the structural optimization, functional validation, and mechanistic analysis of key peptides such as AWPGPQ. On the experimental front, we plan to first complete the chemical synthesis of AWPGPQ [42] and conduct systematic verification of its antioxidant and anti-muscle atrophy effects in both cell and animal models to clarify its biological activity and stability in vitro and in vivo. For structural optimization, we will explore strategies such as D-amino acid substitutions [43] and N-terminal modifications [44] proposed by Jia, Li, and others to enhance its enzymatic stability and in vivo stability, thus improving its drug potential. Mechanistic studies will consider introducing CRISPR/Cas9 technology [45], as applied by Matsunaga *et al*., to knock down key genes of *Keap1* and *PPARγ*, combined with fluorescence reporter systems to analyze the regulatory effects of AWPGPQ on downstream signaling nodes (such as *Nrf2* nuclear translocation and *PPARγ* transcriptional activity), further validating its dual-pathway synergistic mechanism. Provided sufficient technical and resource support, future studies will also investigate its feasibility for application in functional foods or natural therapeutics, gradually advancing the translation of basic research findings into practical applications, thus contributing to the high-value utilization and scientific development of marine active peptides.

## 4. Materials and Methods

### 4.1 Preparation and Preliminary Physicochemical Characterization of Oyster Peptides

The oyster samples from Rushan were soaked in 0.1 mol/L NaOH for 30 minutes, then washed with distilled water until the pH reached 9. Alkaline protease (6000 U/g) was used for enzymatic hydrolysis at 50°C with stirring (400 rpm) for 4 hours [46]. After the hydrolysis reaction was terminated, the enzyme was inactivated by boiling for 10 minutes, followed by centrifugation at 4000 rpm for 15 minutes at room temperature. The supernatant was collected and freeze-dried to obtain oyster peptides. The protein and moisture contents were determined according to the standards GB 5009.5-2016 and GB 5009.3-2016, respectively. The molecular weight distribution was determined according to GB/T 22729-2008, using a mobile phase consisting of acetonitrile, water, and trifluoroacetic acid in a ratio of 40:60:0.2 (v/v/v), with a standard molecular weight solution concentration of 0.2 mg/mL.

### 4.2. Establishment of C2C12 Muscle Atrophy Model and In Vitro Activity Evaluation

Mouse C2C12 myoblasts were obtained from Beijing Bena Chuanglian Biotechnology Institute and cultured in DMEM medium (C11995500BT, Gibco, USA) containing 10% FBS (Zhongke Maichen, China) at 37°C in a 5% CO_2_ incubator. Differentiation was induced by culturing the cells in DMEM medium with 2% horse serum for 7 days [47], with media replaced every 2 days. During the logarithmic growth phase, C2C12 cells were seeded into 96-well plates (100 μL/well) and cultured for 24 hours until confluence reached 70–80%, then the medium was replaced with serum-free DMEM and treated with different concentrations of peptides: oyster peptides were used at 25, 50, 100, 200 μg/mL, and DEX at 5, 10, 20, 40 μg/mL. A control group was also set up, with five replicates per group. After 24 hours of incubation, 90 μL of serum-free DMEM and 10 μL of MTT solution (5 mg/mL) were added to each well [48] and incubated for 4 hours. After discarding the supernatant, 100 μL of DMSO was added and shaken for 10 minutes. Cell viability was analyzed using a CCK-8 assay kit for both experimental and control groups [49].

### 4.3. Functional Indexes of Muscle Atrophy Regulated by Oyster Peptides

C2C12 cells were seeded in 6-well plates and washed three times with PBS, followed by fixation with 4% paraformaldehyde (White Shark, China) for 10 minutes. The cells were treated with 0.3% Triton X-100 (Lamboleid, China) for 15 minutes to enhance membrane permeability [50], followed by blocking with 3% BSA for 2 hours. The cells were incubated overnight at 4°C with a 1:100 dilution of MyHC primary antibody [51], and the next day, secondary antibody was added, followed by nuclear staining with DAPI (Lamboleid, China). Fluorescent images were captured using a fluorescence microscope, and MyHC-positive nuclei, fusion index, myotube number, width, and MyHC-positive area were analyzed using ImageJ software. Data were obtained from at least three independent cultures.

For the 25 and 50 μg/mL oyster peptide-treated cells, total RNA was extracted using the Trizol method, and after determining purity and concentration, reverse transcription was performed using the Takara reverse transcription kit to generate cDNA. qRT-PCR was conducted using SYBR Green Premix [52] on the Roche LightCycler 480 system, with primers designed for *Atrogin-1, MuRF1*, and *GAPDH* as the reference gene. The nucleotide sequences of the primers are shown in Table 4. The relative expression levels were calculated using the ΔΔCT method, with three replicates per group, and the results were used for gene expression analysis.

**Table 4.**
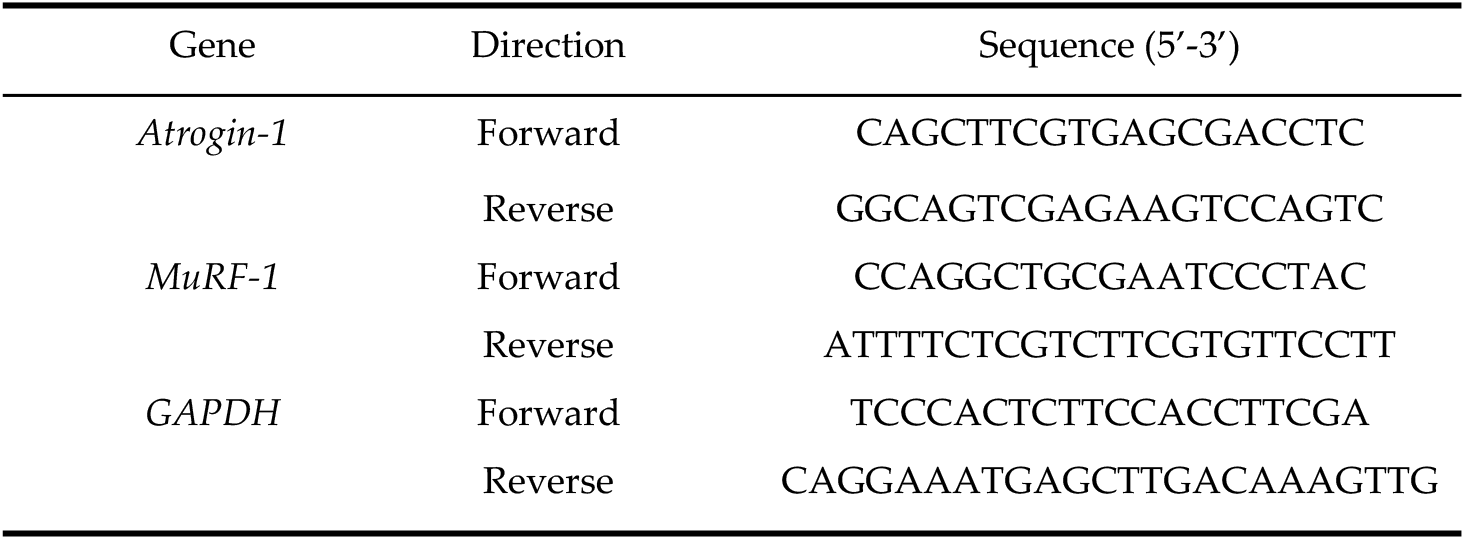
Primers Used for Quantitative Real-Time PCR.

### 4.4. Isolation, Sequencing, and Activity Screening of Active Components in Oyster Peptides

The crude oyster peptides were dissolved in 0.01 mol/L citrate-citrate buffer (200 mg/mL), filtered through a 0.45 μm membrane, and 1.0 mL of the sample was loaded onto a Sephadex G-25 gel column (2.0 × 120 cm) [53]. The elution was performed at a flow rate of 2 mL/min, and one tube was collected every 2 minutes, totaling 100 tubes. Absorbance was monitored at 220 nm to obtain the elution profile. The main peaks were collected and freeze-dried, yielding the fractions OP1, OP2, and OP3. These three fractions were tested for their effects on C2C12 cell viability and their regulation of muscle atrophy marker gene expression.

The oyster peptide samples were dissolved in ultrapure water, treated with DTT (56°C for 1 hour) and IAA (in the dark for 40 minutes), and then centrifuged at 4000 rpm using a high-speed centrifuge (Fresco 17, Thermo Fisher Scientific, China) to remove the supernatant and desalt. The samples were then vacuum-dried using a vacuum concentrator (CV100-DNA, Beijing Jiayim Technology Co., Ltd.). Next, peptides were separated using a high-performance liquid chromatography system (Easy-nLC 1200, Thermo Scientific) equipped with an Acclaim PepMap C18 column (150 μm × 150 mm, 1.9 μm), using 0.1% formic acid (FA) as mobile phase A and 0.1% FA + 80% acetonitrile (ACN) as mobile phase B, with gradient elution [54]. The separated products were identified using an Orbitrap Fusion Lumos high-resolution mass spectrometer (Thermo Scientific) in combination with LC-MS/MS technology [55], and the data were analyzed using PEAKS software. The false discovery rate (FDR) was used to evaluate data reliability, and peptides with a De novo score >90% were selected as high-confidence peptides [56]. The biological activity potential of these peptides was predicted using the PeptideRanker server (http://distilldeep.ucd.ie/PeptideRanker/) and peptides with a predicted biological activity score ≥0.7 were selected for further functional studies.

### 4.5. Construction and Evaluation of Machine Learning Models for Antioxidant Peptide Prediction

This study replicated and optimized the antioxidant peptide prediction model (iAnOxPep) by Hassan *et al*. [20], adding SVM and MLP classifiers to the original model and extending the feature encoding to 98 dimensions. The core structure of the original model was inherited, and seven complementary feature encodings [20] were used to construct a multi-dimensional peptide sequence representation: amino acid composition (AAC, 20 dimensions), adjacent dipeptide frequency (DPC, 400 dimensions), sequence order-dependent features (*PC-PAAC* and *SC-PAAC*, 22 dimensions each), multi-scale structural features (CTD, 147 dimensions), K-gap dipeptide composition (KSDC, 800 dimensions), and spatial distribution features (PDT, 531 dimensions) [57]. Seven base classifiers were trained under these seven features, including random forest (RF), XGBoost (XGB), LightGBM (LGBM), CatBoost, Extra Trees Classifier (ETC), and the newly added support vector machine (SVM) and multi-layer perceptron (MLP), generating 49 prediction probabilities (PP) and 49 classification labels (CL), forming a 98-dimensional high-discriminatory meta-feature vector. This vector was input into five ensemble models (RF, XGB, LGBM, ETC, CatBoost) for weighted voting to output the final prediction results.

In terms of data, we integrated PeptideAtlas and AODP databases based on APD3, BIOPEP-UWM, and PEPLAB, removing sequences with non-standard amino acids or abnormal lengths and using BLAST to eliminate redundant sequences with a similarity greater than 95%. A total of 2,343 positive samples and 7,268 negative samples were retained. For data preprocessing, StandardScaler was used for feature normalization [59], and stratified K-fold cross-validation was employed to maintain consistent class distribution. During training, Optuna was used for Bayesian hyperparameter optimization, and 5-fold cross-validation was applied to enhance model stability [60]. Isotonic regression based on cross-validation was used to calibrate the prediction probabilities, improving reliability, and GPU acceleration was enabled for classifiers like XGBoost, LightGBM, and CatBoost to optimize training efficiency.

Finally, the model was comprehensively evaluated using five metrics: accuracy, specificity, sensitivity, AUC, and MCC [20,61], which measured overall prediction accuracy, the ability to discriminate negative samples, positive sample recognition, model stability under varying classification thresholds, and the correlation between predicted and actual outcomes [62].

### 4.6. Molecular Docking Validation of Antioxidant and Anti-Atrophy Peptides

#### 4.6.1. Preparation of Docking Files

The targets used for molecular docking were *Keap1* (PDB ID: 5FZN) and *PPARγ* (PDB ID: 1FM6), with their three-dimensional crystal structures obtained from the RCSB Protein Data Bank. Protein structures were preprocessed using PyMOL software [63], removing water molecules, non-essential ligands, and heteroatoms, and adding polar hydrogen atoms to optimize the structure [64]. The processed proteins were converted into PDBQT format using AutoDock Tools 1.5.6 [65] for docking with AutoDock Vina.

The peptide ligands were drawn in 2D using ChemDraw software [66], and their correct spatial orientation was assigned. The 2D structures were then converted to 3D conformations using Chem3D, and energy minimization was performed to ensure their stability [67], providing suitable starting structures for molecular docking.

#### 4.6.2. Molecular Docking Simulation

Molecular docking was performed using AutoDock Vina software, which utilizes a gradient-optimized heuristic global-local hybrid search algorithm [68] to predict the binding conformations of ligands and receptors. During the docking process, the active site of *Keap1* was defined with a docking box centered at coordinates (center_x = 13.7 Å, center_y = 65 Å, center_z = 30 Å), with box dimensions of (size_x = 14 Å, size_y = 29 Å, size_z = 51 Å). For *PPARγ*, the active site was defined with a docking box centered at (center_x = 18.7 Å, center_y = −19.2 Å, center_z = 2 Å), and dimensions of (size_x = 58 Å, size_y = 63.2 Å, size_z = 71.8 Å). A semi-flexible docking strategy was employed (ligand flexibility with rigid receptors), and batch scripts were used to generate 9 optimal binding conformations for each ligand-receptor system, with detailed docking reports generated.

The AutoDock Vina scoring function evaluates the binding affinity of ligands to receptors based on factors such as van der Waals forces, electrostatic forces, hydrogen bonding, and desolvation energy [68,69]. According to the research by Guan [70], Liu [71], and others, the active peptide sequence YYR and rosiglitazone (RSG) are known ligands of *Keap1* and *PPARγ*, respectively, and exhibit anti-muscle atrophy activity [70,71]. These were set as control groups and docked with their respective receptors. Meanwhile, the peptides selected through machine learning were also docked with the aforementioned receptors, and their interaction strength was evaluated based on the binding energy. The binding energy data were normalized by subtracting the mean and dividing by the standard deviation to calculate the Z-score for each peptide sequence [72], which quantifies its performance relative to the overall predictions. The Z-scores were then linearly transformed into the [0,1] range and sorted from best to worst.

#### 4.6.3. Visualization and Analysis of Docking Results

Based on the Z-score ranking, the top-quality peptide sequences were analyzed using PyMOL [73] and LigPlus software to examine the docking results and conformations, focusing on key amino acid residues, hydrogen bond interactions, and hydrophobic interaction patterns [74]. Based on these analyses, the binding abilities of the peptides with *Keap1* and *PPARγ* were assessed.

### 4.7. Molecular Dynamics Simulation to Validate the Stability of Peptide-Target Binding

An 80 ns molecular dynamics simulation was performed on the ligand-receptor complexes [75,76]. Simulations were carried out using Gromacs 2023.2 software with the CHARMM36 force field for proteins [77], the AMBER GAFF force field for ligands, and the TIP3P water model [58]. The system was placed in a cubic water box with a side length of 16.118 nm [78], and ions were added to neutralize the system’s net charge. During energy minimization, the first 10,000 steps used the steepest descent method, followed by 10,000 steps of conjugate gradient optimization, with a convergence criterion of energy change less than 1000 kJ/mol. In the equilibration phase, 100,000 steps were performed under both NVT and NPT ensembles to achieve the target temperature, pressure, and density.

The stability and interactions of the complexes were assessed through multiple indicators: RMSD (Root Mean Square Deviation) was used to evaluate the overall structural stability of the complex [79]; RMSF (Root Mean Square Fluctuation) quantified the fluctuations of individual amino acid residues [80]; Rg (radius of gyration) measured the compactness of the system; the number of hydrogen bonds formed between the ligand and receptor reflected the stability of the binding interface. Additionally, the Gibbs free energy of ligand-receptor binding was calculated using Python scripts [81], and the free energy landscape was visualized by plotting the RMSD and Rg values [82]. The lowest free energy dominant conformational clusters were identified, and the key strong interactions within these stable conformations were analyzed [82,83].

### 4.8. ADMET Property Prediction and Drugability Evaluation

To verify the drug potential of the selected active peptides, we used the admetSAR network server (https://lmmd.ecust.edu.cn/admetsar2/) to evaluate their absorption, distribution, metabolism, excretion, and toxicity (ADMET) properties by inputting the molecular structure of the peptides [84], providing theoretical support for subsequent drug development.

### 4.9. Experimental Data Statistical and Analytical Methods

In this study, all experimental data are presented as mean ± standard error of the mean (Mean ± SEM), and statistical analysis was performed using SPSS 20.0 software. Differences in cell viability and gene expression levels were assessed using one-way analysis of variance (ANOVA). Tukey HSD multiple comparisons were used when the data met normality and homogeneity of variance assumptions; otherwise, the Kruskal-Wallis H test was applied. The significance of differences between groups was assessed using Dunnett’s test. Statistical significance was set as follows: *P* < 0.05 was considered significant (*), *P* < 0.01 was highly significant (**), and *P* < 0.001 was very highly significant (***).

## 5. Conclusions

This study systematically evaluated the intervention effects of oyster peptides on the DEX-induced C2C12 muscle atrophy model. The results demonstrated that oyster peptides, within a concentration range of 25–50 μg/mL, significantly improved cell viability, alleviated muscle fiber atrophy, and effectively downregulated the expression of atrophy marker genes *atrogin-1* and *MuRF-1*, indicating strong anti-atrophy potential. The main active component, OP-3, was obtained through gel filtration chromatography and combined with the identification of 220 high-confidence peptide segments using LC-MS/MS. Through an integrated machine learning model, 15 candidate antioxidant peptides were selected. Subsequent molecular docking and dynamics simulations validated the strong binding affinity structural stability of AWPGPQ with both key targets, Keap1 and PPARγ, suggesting that AWPGPQ may regulate the *Keap1*-*Nrf2* and *PPARγ* pathways to collaboratively intervene in the muscle atrophy process. ADMET analysis further supported that AWPGPQ has good safety and drug-like potential. However, the study requires functional validation of the synthesized peptide to confirm its in vivo activity and further verification of its mechanism and applicability across multiple species models. Future research will focus on in-depth mechanism analysis and peptide structure optimization to enhance its translational application in new drug development. In conclusion, our research not only provides a high-efficiency strategy for screening active peptides but also offers valuable insights for the development of multifunctional peptides to combat muscle atrophy.

## Supplementary Materials

The following supporting information can be downloaded at: https://zenodo.org/records/16436371, Figure S1: False Discovery Rate (FDR) Curve; Figure S2: PEAKS Score Distribution and Mass Error; Figure S3: Precursor Mass Error of PSMs; Figure S4: Binding Mode of Keap1 with FGPF and NFNF; Figure S5: Binding Mode of PPARγ with FGPF and NFNF; Figure S6: Solvent-Accessible Surface Area (SASA) Visualization of AWPGPQ Bound to Keap1 and PPARγ; Table S1: Peptide and Protein Analysis Overview; Table S2: Result Filtration Parameters and Filtered Results Statistics; Sheet 1: LC-MS/MS Analysis Data; Sheet 2: PeptideRanker Bioactivity Scores; Sheet 3: Machine Learning Predictions and Probabilities.

## Author Contributions

Conceptualization, J.Z. (Jiyuan Zhang), L.L. (Linlun Zhou) and T.F.(Ting Fang); methodology, J.Z. and L.L.; software, J.Z. and J.Y. (Jiawei Yang); validation, J.Z., L.L., J.Y. and H.Z. (Hailin Zheng); formal analysis, J.Z., L.L. and J.Y.; investigation, J.Z., L.L. and J.Y.; resources, J.Z., L.L., J.Y. and H.Z.; data curation, J.Z. and J.Y.; writing—original draft preparation, J.Z. and L.L.; writing—review and editing, J.Z and W.J. (Wenjuan Li); visualization, J.Z., L.L. and J.Y.; supervision, J.Z. and W.L.; project administration, J.Z., L.L., T.F and W.L.; funding acquisition, J.Z. and W.L. All authors have read and agreed to the published version of the manuscript.

## Institutional Review Board Statement

Not applicable.

## Data Availability Statement

This study utilized publicly available datasets, as cited in the manuscript. All other data generated or analyzed during the current study are included in this published article.

## Abbreviations

The following abbreviations are used in this manuscript:

AAC: Amino Acid Composition
CAN: Acetonitrile
ADMET: Absorption, Distribution, Metabolism, Excretion and Toxicity Atrogin-1 Atrophy-related gene 1
BCA: Bicinchoninic Acid
BBB: Blood-Brain Barrier
CTD: Composition, Transition, Distribution CYP450 Cytochrome P450 Enzyme
DAPI: 4′,6-diamidino-2-phenylindole DEX Dexamethasone
DMEM: Dulbecco’s Modified Eagle Medium DMSO Dimethyl Sulfoxide
DPC: Dipeptide Composition
DTT: Dithiothreitol
ETC: Extra Trees Classifier
FA: Formic Acid
FBS: Fetal Bovine Serum
FDR: False Discovery Rate
GAPDH: Glyceraldehyde-3-Phosphate Dehydrogenase HIA Human Intestinal Absorption
HPLC: High Performance Liquid Chromatography IAA Iodoacetamide
iAnOxPep: integrated Antioxidant Peptide Prediction model Keap1 Kelch-like ECH-associated protein 1
LGBM: Light Gradient Boosting Machine
LC-MS/MS: Liquid Chromatography–Tandem Mass Spectrometry MLP Multi-Layer Perceptron
MuRF1: Muscle RING-finger protein-1 MyHC Myosin Heavy Chain
MTT: 3-(4,5-Dimethylthiazol-2-yl)-2,5-diphenyltetrazolium bromide
PPARγ: Peroxisome Proliferator-Activated Receptor Gamma
PBS: Phosphate-Buffered Saline
PC-PAAC: Position-Specific Physicochemical Properties–Pseudo Amino Acid Composition
PDT: Pseudo Distance Transformation
qRT-PCR: Quantitative Real-Time PCR Rg Radius of Gyration
RF: Random Forest
RMSD: Root Mean Square Deviation
RMSF: Root Mean Square Fluctuation ROS Reactive Oxygen Species
SC-PAAC: Sequence-Order-Coupling Number–Pseudo Amino Acid Composition
SDS-PAGE: Sodium Dodecyl Sulfate–Polyacrylamide Gel Electrophoresis
SVM: Support Vector Machine
UV: Ultraviolet
XGB: Extreme Gradient Boosting hERG human Ether-à-go-go-Related Gene

