## Supplement table1 for "Screening of Oyster Peptides for Anti-Muscle Atrophy Based on Machine Learning and Computer Simulation: Guided by Antioxidant Pathways": Supplementary materials.docx

**Table S1.** Peptide and Protein Analysis Overview

|  | #Scans | | | #Features | Identified | | | #Peptides | #Sequences | #Proteins^*^ | | |
| --- | --- | --- | --- | --- | --- | --- | --- | --- | --- | --- | --- | --- |
|  | MS1 | MS/MS | #Chimera |  | #PSMs | #Scans | #Features^**^ |  |  | Groups | All | Top |
| Total | 9630 | 37280 | 9109 | 44310 | 62 | 62 | 49 | 33 | 31 | 9 | 38 | 14 |

**Table S2.** Result Filtration Parameters and Filtered Results Statistics

| Peptide -10lgP | ≥ 18.29 | FDR (Peptide-Spectrum Matches) | 1.00% |
| --- | --- | --- | --- |
| Protein -10lgP | ≥ 20 | FDR (Peptide Sequences) | 0.00% |
| Proteins unique peptides | ≥ 1 | FDR (Protein Group) | 0.00% |
| De novo score(%) | ≥ 90% | De Novo Only Spectra | 603 |

**Figure S1.** False Discovery Rate (FDR) Curve
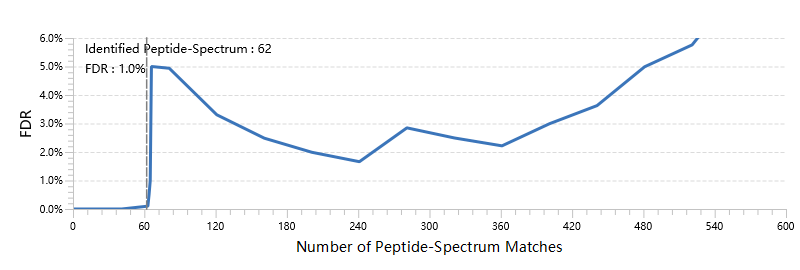

**Figure S2.** PEAKS Score Distribution and Mass Error
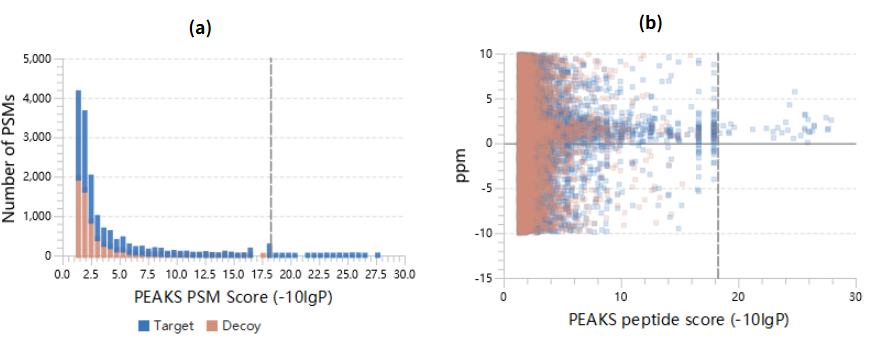

**Figure S3.** Precursor mass error of peptide-spectrum matches (PSM) in filtered result. **(a)** Distribution of precursor mass error in ppm; **(b)** Scatterplot of precursor m/z versus precursor mass error in ppm.

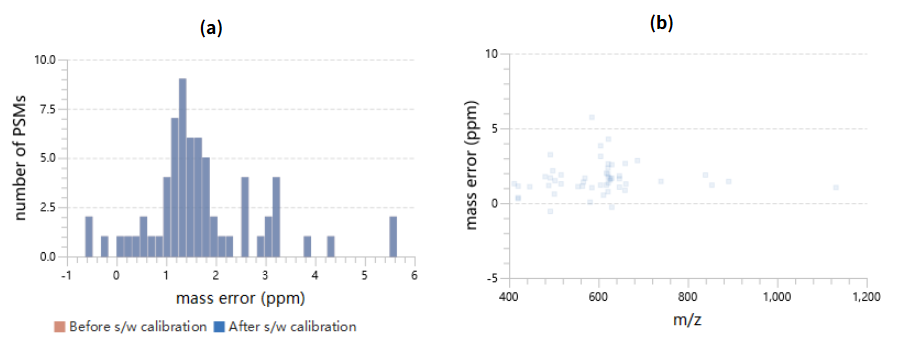

**Figure S4.** Binding Mode of Keap1 with FGPF and NFNF

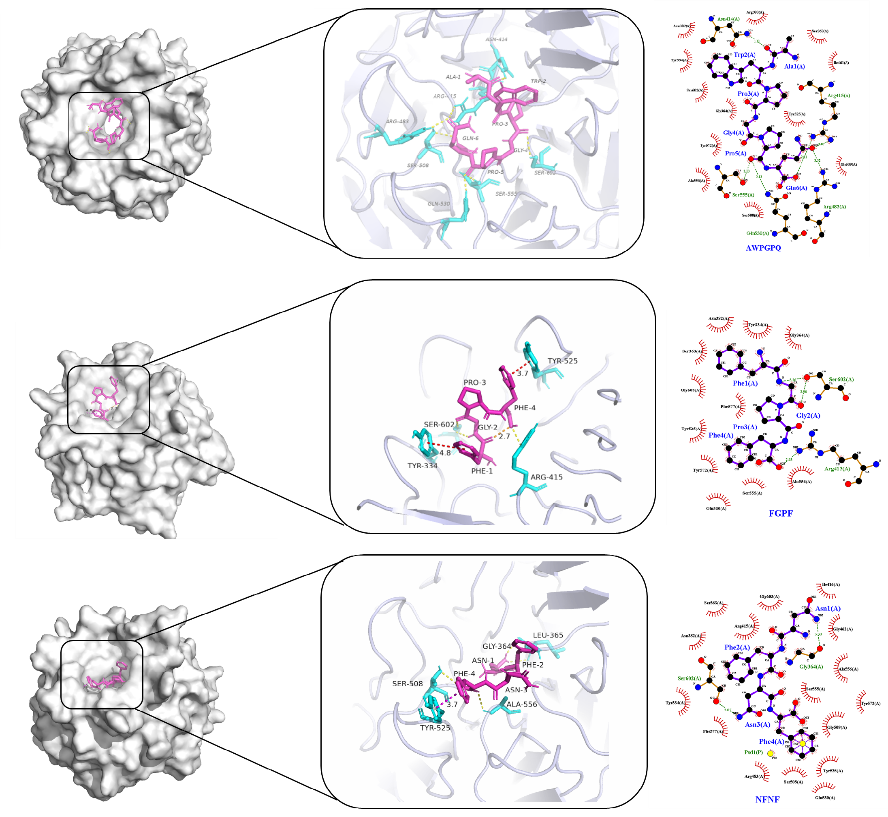

**Figure S5.** Binding Mode of PPARγ with FGPF and NFNF

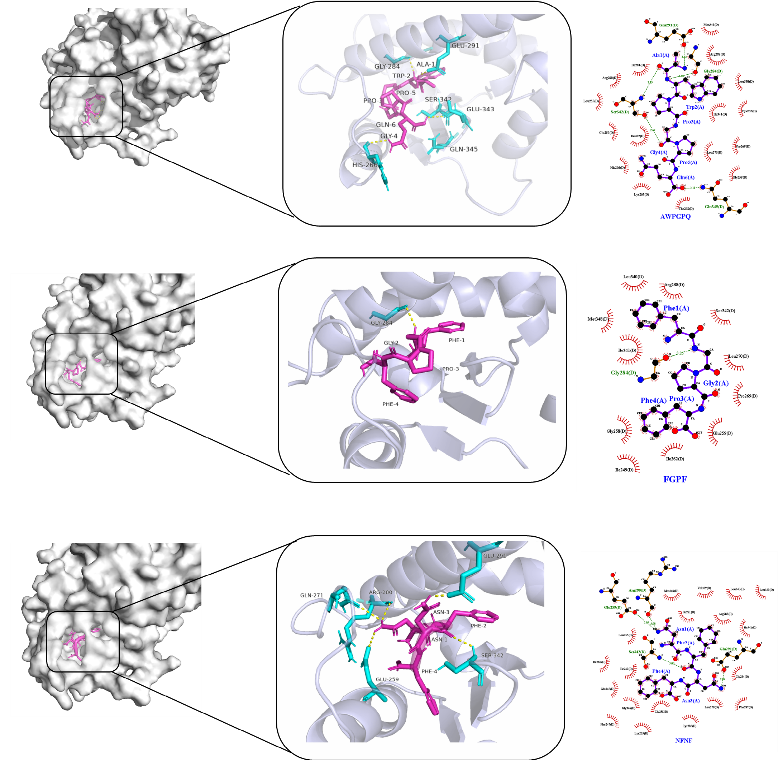

**Figure S6.** Solvent-accessible surface visualization of AWPGPQ bound to (a) Keap1 and (b) PPARγ, showing binding interfaces and surface topologies.

(b)

(a）

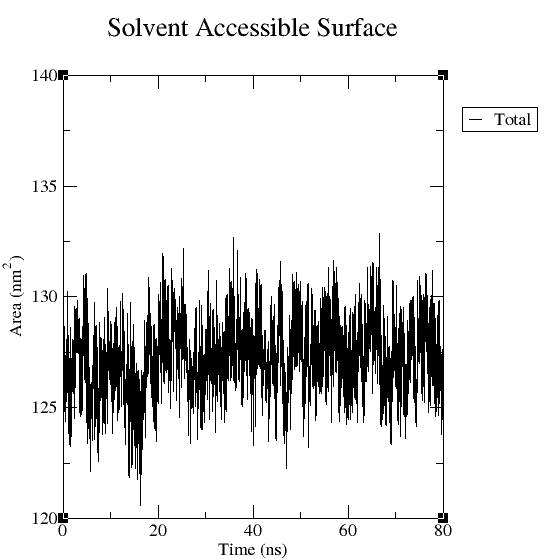

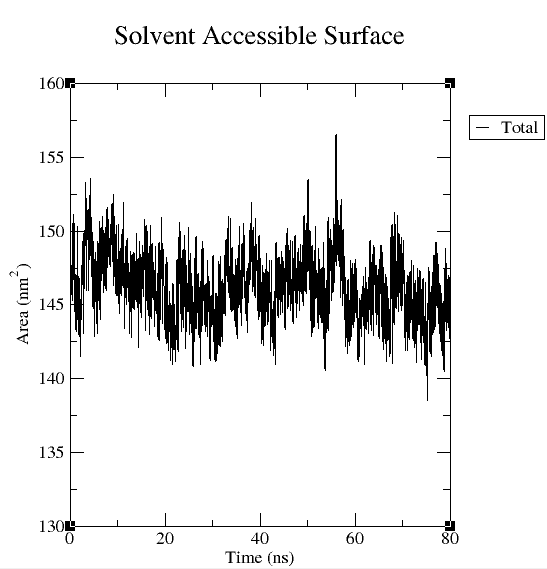
